# Heterogeneous neuronal activity in the ventral tegmental area coordinates dopamine release in the nucleus accumbens

**DOI:** 10.1101/2022.07.31.502221

**Authors:** Daniel F. Hill, Zachary G. Olson, Mitchell J. Bartlett, Torsten Falk, Michael L. Heien, Stephen L. Cowen

## Abstract

Dopamine release in the ventral striatum is fundamental to adaptive appetitive behavior. Frustratingly, technological, and methodological hurdles have limited the investigation of the mechanisms that drive phasic dopamine release. Although dopamine neuron activation in the ventral tegmental area (VTA) surely results in dopamine release, how populations of these neurons coordinate their activities in time to modulate the temporal pattern of release remains unclear. Additionally, burgeoning evidence suggests that control over striatal dopamine release is not solely regulated by the burst firing of VTA neurons or even cell-body activation of these neurons. Here, we recorded VTA neurons while simultaneously monitoring pharmacologically induced phasic dopamine release events in the ventral striatum (nucleus accumbens core) of anesthetized rats. On average, dopaminergic and non-dopaminergic neurons increased activity at the time of the onset of release and decreased at the time of peak release; however, the tuning of individual VTA neurons to dopamine release was notably heterogenous, with subsets of neurons responding prior to release, at release onset, or during release. Other neurons were notably silent during release but active otherwise. Interestingly, both putative dopaminergic and non-dopaminergic neurons expressed this temporally heterogeneous response pattern. Furthermore, the firing activity of dopaminergic, but not non-dopaminergic neurons, correlated with the magnitude of dopamine release. These data suggest that populations of VTA neurons become active at distinct times of a dopamine release event to sculpt the temporal pattern of release.

## Introduction

Dopaminergic neuron activity in the ventral tegmental area (VTA) and dopamine (DA) release in the nucleus accumbens (NAc) are essential for decision making (Schultz, 2015; Schultz et al., 2017), error-driven learning (Eshel et al., 2016; Schultz et al., 2015), and addiction (Lüscher, 2016). While it is well established that increased dopaminergic neuron activity in the VTA is associated with increased dopamine release in downstream structures (Bass et al., 2010; Lohani et al., 2018; Melchior et al., 2015; Sombers et al., 2009), this relationship has never been directly investigated at the timescale of action potentials. This is partly due to the limited availability of instrumentation for the simultaneous measurement of fast, “phasic”, changes in dopamine concentration ([DA]) and single-neuron firing activity (Kuhr et al., 1987; Parent et al., 2017). Consequently, little is known about the precise temporal relationships between midbrain dopamine and non-dopamine neuron activity and downstream dopamine release in the NAc.

The time course of dopamine release (*e*.*g*., release rate and duration) can selectively affect learning (Hart et al., 2014; Schultz et al., 2015; Stefani and Moghaddam, 2006), decision making (Sugam et al., 2012), and memory formation (Lisman and Grace, 2005). Indeed, there is evidence that the early, fast, stages of release (< 100 ms) encode stimulus salience (Kobayashi and Schultz, 2014; Redgrave and Gurney, 2006), while release at later stages (100 – 200 ms) correlates with reward prediction errors (Schultz, 2016a). It is also clear that the time course of release can be a complex and non-monotonic function of input to the VTA and NAc (Hill et al., 2017; Jackson et al., 2001). Therefore, it is conceivable that subpopulations of VTA neurons are active at unique phases of each dopamine release event to shape the time course of release.

Interactions between dopaminergic and non-dopaminergic VTA neurons likely control the temporal pattern of dopamine release. As many as 50% of VTA neurons are non-dopaminergic (Margolis et al., 2006), and VTA glutamatergic neurons can respond to rewards and aversive stimuli (Root et al., 2018). VTA gamma-aminobutyric acid (GABA) neurons may also play key roles in regulating the time course of release through disinhibition of dopaminergic neurons (Bourdy and Barrot, 2012; Jalabert et al., 2011; Jhou et al., 2009; Morales and Margolis, 2017; Omelchenko and Sesack, 2009). Such inhibitory control may also trigger bursts and pauses in firing activity in VTA dopaminergic neurons (Paladini and Roeper, 2014).

Dopaminergic neurons themselves can be divided into subsets that respond to rewards (Mirenowicz and Schultz, 1996; Schultz et al., 2015, 1993), aversive stimuli (Badrinarayan et al., 2012; Brischoux et al., 2009; de Jong et al., 2019; Matsumoto and Hikosaka, 2009; Menegas et al., 2018), salience (Fiorillo et al., 2013; Kobayashi and Schultz, 2014; Schultz, 2016b), body movements (Da Silva et al., 2018; Engelhard et al., 2019; Howe and Dombeck, 2016; Ogasawara et al., 2018), and working memory (Matsumoto and Takada, 2013). The diverse functional and temporal responses of VTA neurons reviewed above supports the hypothesis that heterogeneous groups of VTA neurons become active under different conditions and time points to shape the downstream pattern of dopamine release. Furthermore, given evidence for local inhibitory regulation of dopamine neuron activity and reward-guided behavior (Matsui and Williams, 2011; Omelchenko and Sesack, 2009; Polter et al., 2018; Tan et al., 2012; van Zessen et al., 2012), we also hypothesized that the activity of putative non-dopaminergic inhibitory neurons will decrease prior to each dopamine release event to disinhibit VTA dopaminergic neurons.

To investigate these hypotheses, we simultaneously measured single-unit activity in the VTA using microelectrode arrays and phasic dopaminergic signaling in the NAc core using fast-scan cyclic voltammetry (FSCV) in anesthetized rats. This was accomplished using a novel technology developed in-house that permits the multiplexing of electrophysiological and voltammetric measurement of dopamine (The DANA system (Parent et al., 2017)). We observed that, on average, dopaminergic and non-dopaminergic neurons increased activity at the time of the onset of release and decreased activity at the time of peak release; however, the tuning of individual VTA neurons to dopamine release was notably heterogenous in time.

## Results

### Identification of dopaminergic and non-dopaminergic neurons

Putative dopaminergic and non-dopaminergic neurons were identified by clustering on half-width and amplitude ratio (**Figure 1C, top**) (Roesch et al., 2007). Neurons near the Fisher’s linear discriminant boundary (indicated with ‘+’) were eliminated to reduce misclassification errors as described in Methods. Each dot in **Figure 1C** indicates an individual neuron, and waveforms from exemplar neurons are presented to the right. Based on these criteria, a total of 105 putative dopamine neurons and 441 non-dopaminergic neurons were identified in 22 rats.

**Figure 1.**
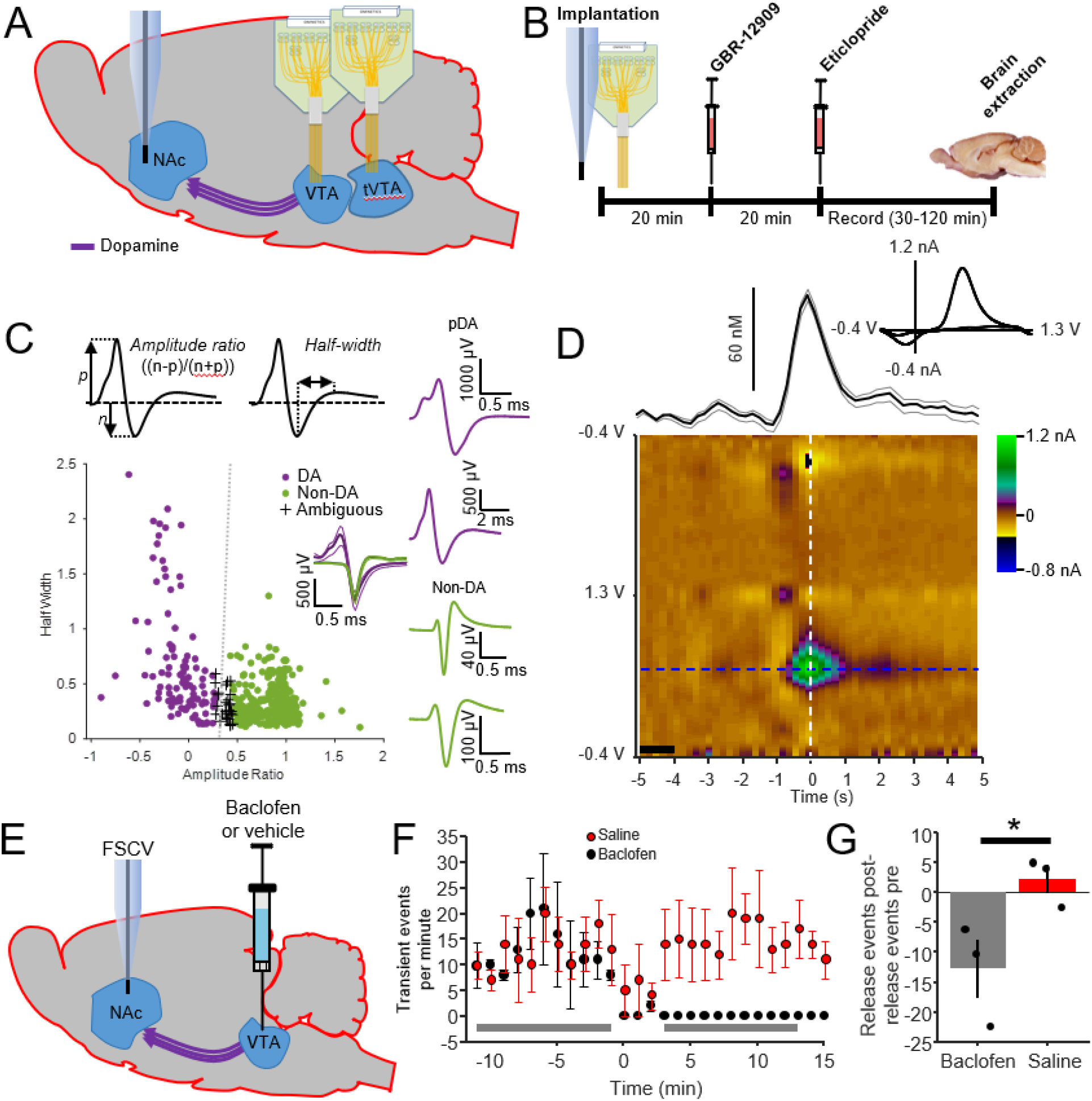
Experimental design. A) Microelectrode arrays (2 × 32 channels) were implanted in into the VTA and tVTA and a carbon-fiber microelectrode was implanted into the NAc. B) After implantation, GBR-12909 and eticlopride were injected to stimulate phasic dopamine release in the NAc. C) Putative dopamine neurons (DA) were identified using clustering based on waveform features (amplitude ratio and half-width). A cell was given membership to a cluster if its half-width and amplitude ratio were within three standard deviations of the centroid based on Euclidean distance (large red and grey circles estimate three standard deviations for all cells in each cluster). Neurons that fell into the region where the large circles overlap were categorized as putative non-dopaminergic. Outliers were removed prior to clustering and categorized individually. The insets above the cluster illustrate the waveform characteristics used to calculate amplitude ration and half-width. Insets to the right show examples of putative dopamine neuron waveforms (red) and non-dopamine neurons (grey). D) Dopamine release events were manually scored to identify onset, offset, and peak. The average of all dopamine release events is shown for one animal. The color plot shows the averaged FSCV signal centered on the peak of all dopamine release events for a single animal. The background was taken from an average of 5 scans from -5 to -4 seconds (n = 258 events). The trace directly above the color plot shows the mean (±SEM) change in dopamine concentration over time and the inset shows a cyclic voltammogram taken at time zero. E) Diagram of recording and injection probes used for the baclofen inactivation of VTA neurons. F) Following injections of GBR-12909 and eticlopride, baclofen was microinjected into the VTA to inhibit dopaminergic neurons. Average of transient dopamine release-event frequency versus time. G) Frequency of pre-baclofen release events (events per minute) was subtracted from post-baclofen release events to measure the change in release following baclofen injection. Release was averaged over the times indicated by the grey bars in subplot F. Administration of baclofen resulted in a significant reduction in spontaneous release events (t-test, p = 0.049, n = 3 rats). Error bars indicate SEM.

### Quantification and verification of pharmacologically evoked dopamine release

Dopamine release was measured using carbon fiber FSCV electrodes lowered into the NAc. Release in anesthetized rats was induced through the delivery of DAT inhibitor GBR-12909 (17.5 mg/kg, *i*.*p*.) and D2R antagonist eticlopride (0.75 mg/kg, *i*.*p*., **Figure 1B**). Dopamine release events were manually scored if they were ≥ 1 nA and were preceded by ≥ 1 second of transient-free baseline. Dopamine release events only emerged following the injection of eticlopride. The mean duration of transient dopamine release events across animals was 4.4 ± 3 s (SD), with an average inter-event interval of 20 ± 33 s (SD). An example of averaged transient release events aligned to peak release is shown in **Figure 1D**. The figure indicates the average across trials (n = 258 events).

### Baclofen inactivation of VTA dopaminergic neurons decreases NAc dopamine release

To verify that dopamine release in the nucleus accumbens resulted from VTA neuron activation, baclofen (GABA_B_ agonist, 0.2 mg in 0.5 µL) or vehicle (0.5 µL saline) was injected into the VTA at 0.5 µL/minute while FSCV was recorded from the NAc (**Figure 1E**). Baclofen reduced the frequency of transient dopamine release events to near zero (**Figure 1F**). The difference score (pre-injection minus post-injection release-event frequency, shown by the grey bars in **Figure 1G**) was significantly lower for baclofen when compared with saline (p = 0.049, Student’s two-sample t-test, n = 3 for each group).

### Responses of individual dopaminergic and non-dopaminergic neurons to the onset and peak of dopamine release

The relationship between VTA single-unit activity and NAc dopamine release was investigated through the simultaneous measurement of NAc dopamine transients and single-unit firing activity. Firing activity was evaluated at the time of transient onset and at the time of peak dopamine concentration (the first peak in the [DA] signal following transient onset). To illustrate, **Figure 2A, B** shows the average change in dopamine concentration from baseline, and **Figure 2C, D** presents the firing activities of two simultaneously recorded VTA neurons aligned to the onset of each NAc dopamine transient release event (n = 420 events). In this example, the putative dopaminergic (**Figure 2C**) and non-dopaminergic (**Figure 2D**) neuron simultaneously responded shortly after the onset of each dopamine transient. This response was short-lived (∼600 ms) relative to the time course of dopamine release in both neurons and terminated before the time of peak dopamine concentration (the peak in **Figure 2B**).

**Figure 2.**
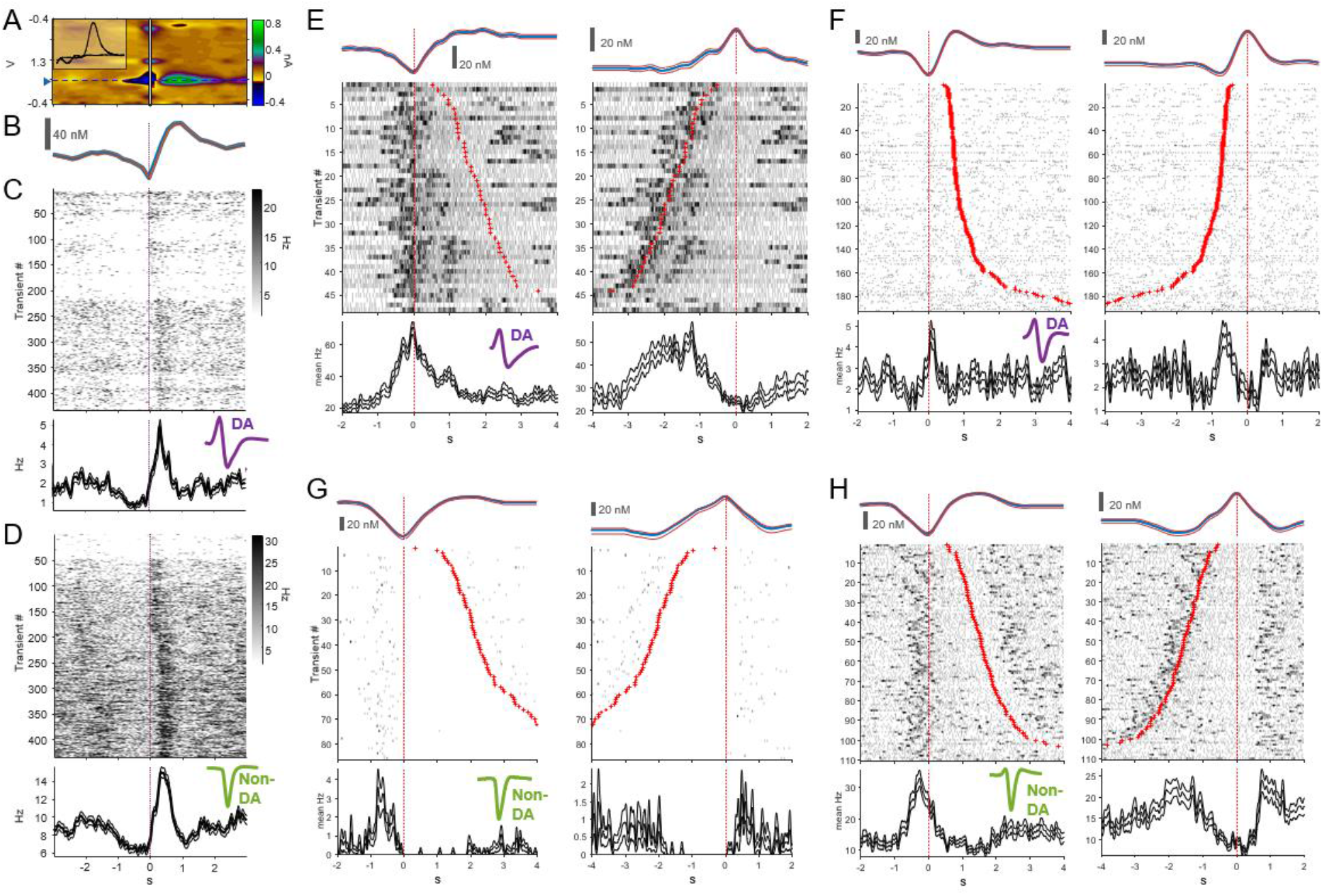
Examples of putative dopaminergic and non-dopaminergic neuronal responses aligned to the onset of dopamine release. A) Average color plot of 420 dopamine release events from a single recording aligned to the scored onsets of the events. Inset depicts the cyclic voltammogram taken from the point of maximal dopamine release. B) Change in dopamine release versus time that corresponds to the blue dashed line and triangle on the color plot in A. C) Response of an exemplar putative dopamine neuron that increases activity following the time of dopamine release onset (time = 0 s). Top: firing response of the neuron surrounding each transient (5 ms bin, smoothed with 40 ms Hanning). Inset: Average waveform shape of the neuron. Below the raster is the mean firing rate (± SEM) versus time the neuron. D) Spike raster and mean response (± SEM) of a simultaneously recorded putative non-dopaminergic neuron that also increased activity following the onset of each dopamine transient. Inset depicts the average waveform of the neuron. E-F) Peri-event responses of two dopaminergic (E,F) and two non-dopaminergic neurons (G,H). As in B, the top trace indicates mean dopamine concentration (± SEM). As in C, individual single-unit responses to each transient are indicated. In these plots, responses to each transient (rows) are sorted according to the duration of each transient, with the transient offset (left panels) or onset (right panels) indicated with a red ‘+’. The left column in each plot indicates the dopaminergic and single-unit response aligned to the transient onset and the right column indicates responses aligned to the time of peak release. The time-aligned responses illustrate the heterogeneity of the single-unit response to the dopamine transients with some neurons responding shortly before (E,G,H) or at the time of onset (F) and others showing a suppression of firing activity at the time of peak release (H).

**Figure 2E-H** illustrate the response of two additional putative dopaminergic (**E, F**) and two non-dopaminergic (**G, H**) neurons to the time of transient onset (left subplots) and the time of peak concentration (right subplots). These examples illustrate the heterogeneity of the single-unit response to the dopamine transients, with some neurons responding shortly before (**E, G, H**) or at the time of onset (**F**) and others showing a suppression of firing activity at the time of peak release (**H**).

### VTA dopaminergic and non-dopaminergic neurons respond near the time of transient onset and peak dopamine release

Given the assumption that VTA neuron activity triggers downstream dopamine release in the NAc, we predicted that VTA neurons would be more active at release onset when compared to the time of peak dopamine release. To test this, neurons that significantly changed their firing activity near the time of transient onset or peak release were first identified. This was determined by first segmenting firing rates into five 800 ms bins centered on release onset or the time of peak release and then performing a within-subject Friedman test to determine if a significant change in activity occurred in one or more of these 800 ms bins (p < 0.05). To validate this criterion, we compared these results to results from a validation set of surrogate single-unit responses generated by shuffling the order of spike counts in the five bins within each trial. Comparison between selective neurons identified to the shuffled control indicated that the Friedman test criterion was effective (See **Supplemental Figure 2**). Neurons were further sub-categorized into cells that increased or decreased firing activity around the onset or peak of release. A neuron was labeled as “increasing” or “decreasing” if the mean firing rate during the significant interval(s) was above or below baseline (one-sample t-test). Baseline was defined as the average activity during the -4 to -2 and +2 to +4 second interval surrounding each release event.

A total of 546 neurons were analyzed (DA: 105, Non-DA: 441). No group differences were observed between the number of dopaminergic and non-dopaminergic neurons responding to release onset (26% DA and 22% of Non-DA neurons) or the peak of release (21% DA and 24% of Non-DA neurons). The mean peri-event responses of all VTA dopaminergic and non-dopaminergic neurons with selective responses are presented in **Figure 3A-D**. Each row in the color plots indicates the response of an individual neuron (color indicates the z score of the firing activity). Neurons were further organized by cells that increased or decreased their activity at release onset (**Figure 3A, B**) or peak release (**Figure 3C, D**). The mean peri-event responses for neurons with increasing (blue) and decreasing (orange) activity are presented at the bottom of each figure, and the increasing and decreasing subsets of neurons are indicated on the y axis.

**Figure 3.**
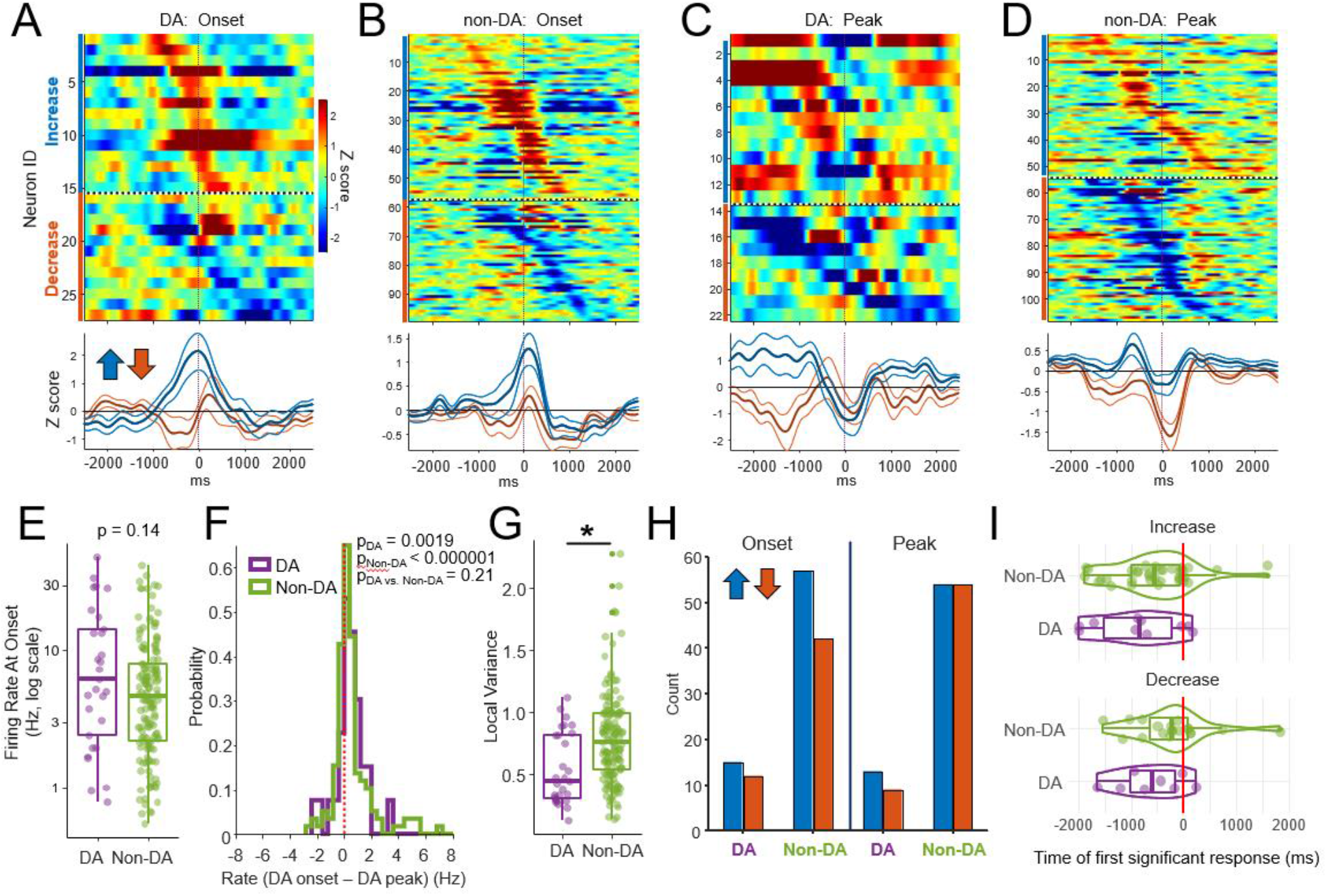
Heterogeneous phasic responses of putative dopaminergic and non-dopaminergic neurons to dopamine release onset and at the peak of release. A-D) Average firing rates of all responsive neurons aligned to phasic dopamine release events. A) Peri-event responses of dopaminergic neurons aligned to the onset of each dopamine transient. Each row indicates the average peri-onset response of a single neuron. Color indicates firing rate in z-scores relative to baseline with baseline defined as activity during the -3 to -4 and +3 to +4 second intervals surrounding the transient onset. The top set of neurons (blue vertical line) increased firing rate activity near transient onset while the bottom set (orange vertical line) decreased activity near transient onset. Line plots below the color plot indicates the mean response (in z scores, +/- SEM) for neurons with responses that increased or decreased firing activity at the time of transient onset. B) As in A, but for non-dopaminergic neurons. C & D) As in A & B, but for neurons selective to the time of peak dopamine release. Baseline was defined as -3 to -4 and +3 to +4 second intervals surrounding the transient peak. E) There was no significant difference in the firing rates of putative dopaminergic and non-dopaminergic neurons during the ± 1 s surrounding release onset (p = 0.14, Wilcoxon). F) Both dopaminergic and non-dopaminergic neurons fired at higher rates at the time of release onset (DA neurons: p = 0.0019, Non-DA neurons p < 0.000001, Wilcoxon signed rank test). This was determined by subtracting the firing rate at the time of peak release from the rate at release onset for each neuron and testing to determine if this distribution of differences was not equal to zero. Firing rates were measured for the -400 to +400 ms interval centered on the onset of release and peak release. There was no significant difference between dopaminergic and non-dopaminergic neurons (p = 0.21, Wilcoxon rank sum test). G) The local-variance measure of inter-spike interval statistics and bursting indicated that the activities of both dopaminergic and non-dopaminergic neurons were decidedly non-bursty (values were, on average, <1). In addition, median local variance was lower in dopaminergic neurons relative to dopaminergic neurons (p = 0.0005 Wilcoxon rank sum test). H) Number of neurons with a significant increase (blue) or decrease (orange) in activity surrounding onset or the peak of release. Comparisons were made between the number of neurons that increased or decreased activity at the time of onset for each cell type (e.g., the number of DA neurons that increased or decreased at the time of transient onset). No difference in the proportion of neurons with increasing or decreasing activity at transient onset or peak were observed (p > 0.05 for all comparisons, binomial test). I) Dopaminergic and non-dopaminergic cells did not differ in the timing of their earliest significant response relative to transient onset. The earliest significant response relative to release onset was determined for each selective neuron (see Methods). While most neurons responded before release onset, there was no difference between the mean timing of release onset between groups (Wilcoxon rank sum test p > 0.05).

Within these groups, neurons (rows) are sorted by the time of the peak or trough of the response. Inspection of the individual single-unit responses indicated considerable heterogeneity in the timing of the maximal and minimal responses relative to the onset or peak of dopamine release. Indeed, mono-phasic and multi-phasic responses at different times relative to the transient onset or peak were observed for both dopaminergic and non-dopaminergic neurons. To better illustrate the heterogeneity of the peri-event responses, we also re-sorted these data using k-means clustered groupings based on the shape of the peri-event responses (**Supplemental Figure 3A-D)**.

Most non-dopaminergic neurons in the VTA are GABAergic (Margolis et al., 2012), and GABAergic neurons can trigger dopamine release by decreasing their activity and thus reducing tonic inhibition of dopamine neurons (Xi and Stein, 1998). Consequently, we predicted that, on average, non-dopaminergic neurons would decrease activity while dopaminergic neurons would increase activity near the time of dopamine release onset. We also predicted that VTA neuron activity would be greatest surrounding release onset relative to the time of peak release as we hypothesized that increased activity prior to or near the time of onset is necessary for initiating phasic release in the NAc. Visual inspection of the mean responses in **Figure 3A-D** and **Supplemental Figure 3** suggested that this was not the case as the mean responses for both DA and non-DA neurons appeared to peak near the time of transient onset. Indeed, the firing rates of putative dopaminergic and non-dopaminergic neurons during the ± 1 s surrounding release onset did not differ between cell types (**Figure 3E**, p = 0.14, Wilcoxon).

Dopaminergic and non-dopaminergic neurons fired at significantly higher rates at the time of transient onset relative to the time of the transient peak (**Figure 3F**, p_DA_ = 0.0019, p_Non DA_ = 0.000001, Wilcoxon signed rank test). Furthermore, no between-group (cell type) difference was detected (p = 0.21, Wilcoxon rank sum test). These results suggest that populations of dopaminergic and non-dopaminergic VTA neurons are involved in the initiation and modulation of dopamine release events, with the caveat that their individual timings were varied around release onset and peak.

### VTA DA and non-DA neurons did not exhibit burst-like firing

Burst firing of dopamine neurons is thought to be a principal driver of dopamine release (Grace and Bunney, 1984). Consequently, we predicted that burst firing should be higher in dopaminergic neurons relative to non-dopaminergic cells and that bursting should increase near the time of measured dopamine release. However, because firing rates of putative dopaminergic neurons were high, likely due to the delivered drugs (*e*.*g*., DAT inhibitor, D2R antagonist), measures of bursting that use rigid inter-spike interval (ISI) thresholds (Grace and Bunney, 1984) would result in many false positives. Thus, a measure of bursting that was robust to differences in firing rate was required. The local variance (LV) measure (Shinomoto et al., 2009, 2005) of bursting and spike-train irregularity addresses this issue and is robust to slow changes in mean firing rates. The LV measure quantifies burst-like responding as the average dissimilarity between adjacent ISIs where more similar intervals suggest tonic/regular firing (LV < 1.0) and more irregular intervals indicate either a Poisson distribution of ISIs (LV = 1.0) or a more long-tailed ISI distribution characteristic of bursting neurons (LV > 1.0). Contrary to our predictions, LV was lower for dopaminergic neurons when compared to non-dopaminergic neurons (**Figure 3G**, p = 0.0005 Wilcoxon rank sum test). Furthermore, the local variance measure did not change from the time of transient onset to the time of peak release (**Supplemental Figure 4**). Thus, at least under these pharmacological conditions (*e*.*g*., isoflurane + GBR-12909 and eticlopride), VTA neural responses did not exhibit burst-like firing.

### Numbers of responsive dopaminergic versus non-dopaminergic neurons did not differ at release onset or peak

Given the hypothesis that transient onset is precipitated by the disinhibition of VTA dopaminergic neurons, we predicted that non-dopaminergic neurons would reduce activity while dopaminergic neurons would increase activity near the time of release onset. Consequently, we investigated whether a larger proportion of non-dopaminergic neurons exhibited a dip in activity (proportionally more ‘decreasing’ responses compared to increasing, **Figure 3H**) relative to dopaminergic neurons. Contrary to this prediction, the proportion of neurons that increased relative to decreased for each cell type did not differ at transient onset (binomial test, p_DA_ = 0.56, p_Non-DA_ = 0.13), or at the time of peak release (p_DA_ = 0.39, p_Non-DA_ = 1.0).

### Both dopaminergic and non-dopaminergic neurons alter their activity ∼1 second before dopamine release onset

We also tested the prediction that decreases in non-dopaminergic neuron activity ought to precede increases in dopamine neuron firing (*e*.*g*., to release dopamine neurons from inhibition). To investigate this question, the time of the earliest significant response for each neuron was quantified as the time when the firing rates in 9 adjacent 15 ms time bins of the peri-event histogram were statistically different from the null distribution. The null distribution was created by randomly shuffling the timing of these 9 bins within each trial (n permutations = 200; significance α = 0.05). While dopaminergic and non-dopaminergic neurons exhibited selective altered activity ∼1 second prior to each identified release event, there was no difference between groups (**Figure 3I**, Wilcoxon rank sum test p > 0.05).

### Varied NAc dopamine dynamics surrounding VTA action potentials

If dopaminergic and non-dopaminergic neurons in the VTA coordinate their activity to initiate dopamine release, it would be expected that the firing activity of VTA neurons would increase when the rate of change of dopamine concentration (Δ[DA]) starts to rise. To investigate this question, we examined the dynamics of dopamine release surrounding VTA action potentials using spike-triggered averages (STAs). STAs and the approximate first derivative of the spike-triggered response (STAs of Δ[DA]) of dopamine concentration were calculated (**Figure 4, Supplementary Figure 5**, see Methods). Briefly, STAs were created by aligning the normalized dopamine concentrations to the time of each action potential, and then averaging these peri-event responses to determine the mean concentrations that surrounded each action potential. To reduce the impact of non-stationarities in the [DA], Δ[DA], and single-unit signals, a surrogate STA was generated by jittering the timing of action potentials (random ±3 s jitter), and this STA was subtracted from the measured STA. A sample of six STAs from dopaminergic and non-dopaminergic neurons to the [DA] and Δ[DA] signals is presented in **Figure 4A, B**. The varied dopamine responses surrounding action potentials in these neurons illustrates diversity in the time course of dopamine release surrounding VTA neuron spiking activity. For example, VTA neurons responded to unique temporal profiles of dopamine release, such as when dopamine levels were rising (**Figure 4A** top row, note positive STA of Δ[DA] at t = 0 s), beginning to rise (**Figure 4A** middle row), or responded to complex multi-phasic features of the dopamine signal (**Figure 4B** top row).

**Figure 4.**
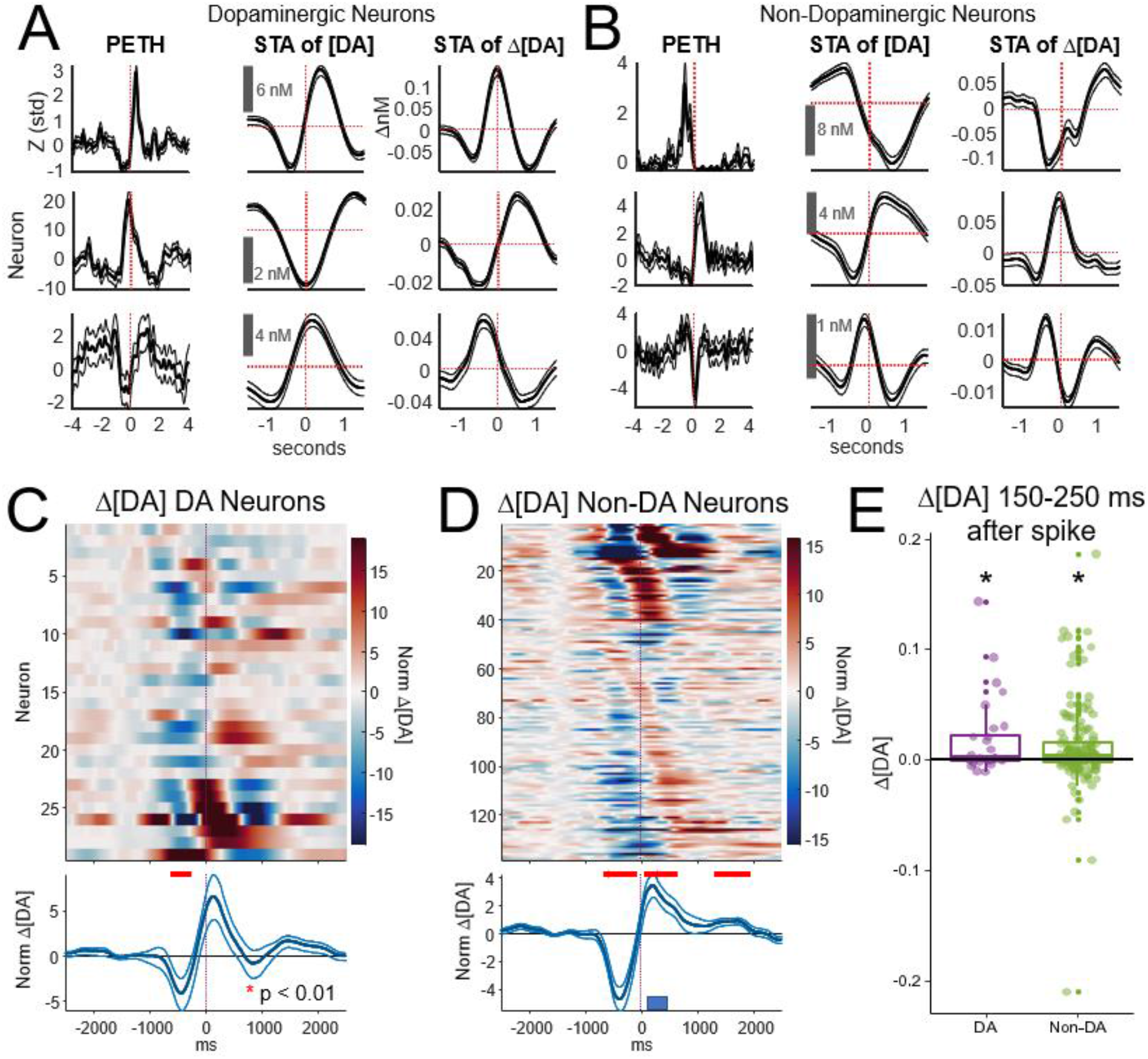
A+B) Mean (±SEM) responses of 3 putative dopaminergic and 3 putative non-dopaminergic neurons. Each row indicates the response of an individual neuron. The first column presents the mean peri-event response (PETH) aligned to the onset of each dopamine release event. Spike-triggered averages (STAs) indicate the mean dopamine concentration aligned to the time of each action potential (vertical line indicates time = 0 s). To reduce the impact of non-stationary changes in dopamine concentrations and single-unit responses, these mean responses were adjusted by subtracting shuffle-averaged STA. The shuffle average STA was created by randomizing the timing of each action potential by ±3 seconds and then using these action potentials to generate a new STA (see Methods). Error bars indicate bootstrapped 95% confidence intervals. The third column shows the change of [DA] (first derivative of data in the STA) and 95% confidence intervals. C+D) Spike-triggered averages (STAs) of the rate of change of dopamine concentration (Δ[DA]) for dopaminergic and non-dopaminergic neurons. C) Each row indicates the STA for a single dopaminergic neuron (see Materials and Methods for details on STA calculation). Color indicates the z-score of Δ[DA] relative to baseline, with baseline defined as activity during the -3 to -4 and +3 to +4 second intervals surrounding t = 0 (the time of each action potential). For visualization and to highlight the diversity of dopamine dynamics surrounding each action potential, STAs are sorted by k-means cluster groups (k = 3) and, secondarily, by the time of the peak response within each group. The bottom panel shows the mean ±SEM. Red asterisks indicate p < 0.01 (Wilcoxon Rank Sum test) relative to zero. D) As in C, but for the non-dopaminergic neurons. E) Mean Δ[DA] was measured for the 150 – 250 ms interval in the mean STA for each neuron. Each point indicates this mean for each neuron. The slope was significantly > 0 for both dopaminergic (p = 0.011, one-sample Wilcoxon test), and non-dopaminergic (p = 0.000015, one-sample Wilcoxon test) neurons. There was no difference between DA and non-DA groups (p = 0.67, 2-sample Wilcoxon test).

Mean STAs of Δ[DA] for all dopaminergic and non-dopaminergic neurons selective to the onset or peak of the transient are presented in **Figure 4C-E**. Each row in **Figure 4C, D** indicates the STA of Δ[DA] for a single neuron. Color indicates the z-score of Δ[DA] relative to baseline, with baseline defined as activity during the -3 to -4 and +3 to +4 second intervals surrounding t = 0 (the time of each action potential) and z = 0 indicating no change in [DA]. The bottom panel shows the mean ± SEM. Horizontal red bars indicate p < 0.01 (Wilcoxon Rank Sum test) to determine if the mean differed significantly from zero. On average, the rate of change of dopamine (Δ[DA]) transitioned from negative to positive surrounding each action potential for both dopaminergic and non-dopaminergic neurons (**Figure 4C, D**). To determine whether this increase followed each action potential on average, mean Δ[DA] was measured for the +150 ms to +250 ms interval following each spike (**Figure 4E**). This time window accounted for the lower sampling rate of the dopamine signal (once every 200 ms). The Δ[DA] signal was significantly > 0 for both dopaminergic (p = 0.011, one-sample Wilcoxon test), and non-dopaminergic (p = 0.000015, one-sample Wilcoxon test) neurons. There was no difference between DA and non-DA groups (p = 0.67, 2-sample Wilcoxon test). This result further supports the hypothesis that VTA spiking from both dopaminergic and non-dopaminergic neurons was positively correlated with downstream dopamine release. It is important to emphasize, however, that while the mean response indicated a positive relationship between VTA neural firing and Δ[DA], there was considerable variation in the responses/STAs of individual neurons. This heterogeneity was also evident in the STAs of the [DA] signal (see **Supplemental Figure 5**).

### Trial-to-trial changes in dopamine-neuron firing activity correlate with trial-to-trial changes in dopamine concentration

If the magnitude of an individual VTA neuron’s firing response determines the magnitude of downstream release, then transient-to-transient changes in [DA] or Δ[DA] should be positively correlated with an individual neuron’s firing activity. To assess this, neural firing rates for a given transient dopamine release event were determined as the mean rate over the -1 to +1 second interval surrounding transient onset. Mean [DA] on a given trial was averaged between +0.15 to +3 seconds following transient onset, and Δ[DA] was averaged between +0.15 and +0.25 seconds following onset (see **Figure 4**). Duration was measured as the time (seconds) from transient onset to peak release. Transient-to-transient dopamine and firing activity measures were then compared using Spearman’s correlation coefficient. The median of the correlation coefficients for DA neurons, but not Non-DA neurons was significantly > 0 (**Figure 5**, p_DA_ = 0.006, p_Non-DA_ = 0.31, sign-rank test, Holm correction for 6 comparisons), and an effect of neuron type (DA vs. Non-DA) was also observed (p = 0.02, Wilcoxson). While population-level activity indicated an overall positive relationship between DA neuron activity and dopamine concentration, the overall proportion of DA and Non-DA neurons that expressed a significant correlation between [DA] and firing activity (p < 0.05) was small (9% of DA neurons, 10% of Non-DA neurons). Correlation coefficients between firing activity and Δ[DA] and duration were not significantly different from 0 (sign-rank-test, p > 0.05). Taken together, and as predicted, these results suggest that trial-to-trial changes in dopamine neuron firing activity (but not Non-DA neuron activity) correlate with downstream release. Interestingly, no relationship was found with Δ[DA] or release duration.

**Figure 5.**
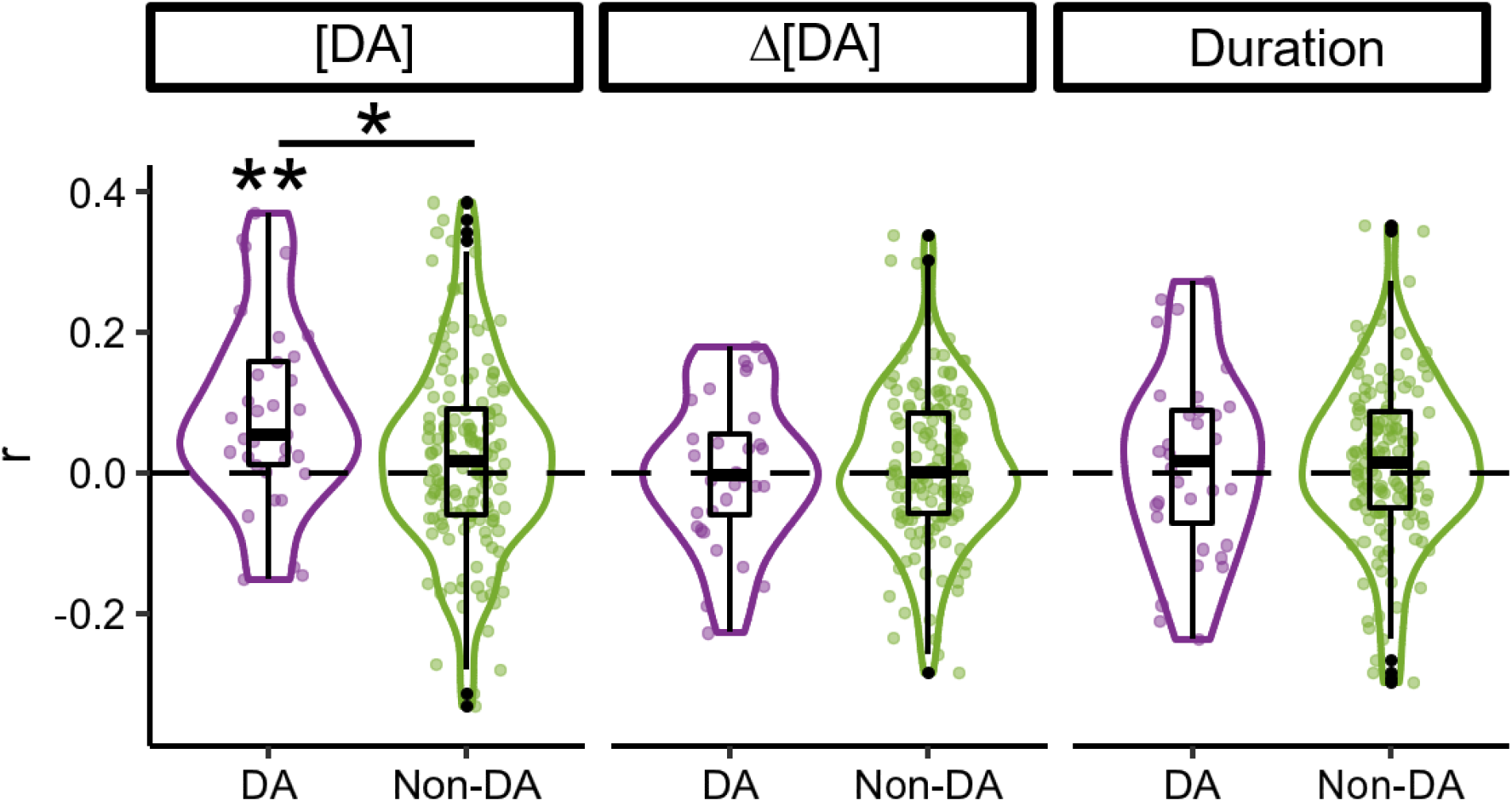
Correlation between trial-to-trial (transient-to-transient) changes in neural firing activity to trial-to-trial measures of dopamine release. Neural firing rates were determined as the mean rate over the -1 to +1 second surrounding transient onset. Mean [DA] on a trial was mean [DA] between +0.15 to +3 seconds following onset, and Δ[DA] was averaged between +0.15 and +0.25 seconds following release (see Figure 5). Baseline activity on each trail was subtracted from each measure to reduce the impact of slow changes in release or firing rate. Duration was the time (seconds) from transient onset to peak release. Trial-to-trial DA and firing activity measures were then compared using Spearman’s correlation coefficient. Baseline-subtracted dopamine concentration ([DA]), the first derivative of dopamine concentration (Δ[DA]), and the duration of each transient were each compared to trial-to-trial firing activity. The median correlation coefficients for DA neurons, but not Non-DA neurons to [DA] was significantly > 0 (**p_DA_ = 0.006, p_Non-DA_ = 0.31, sign-rank test, Holm correction for 6 comparisons), and an effect of neuron type was also observed (*p = 0.02, Wilcoxson). While population-level activity indicated an overall positive relationship between DA neuron activity and [DA] release amount, overall proportion of DA and Non-DA neurons that expressed a significant correlation between [DA] and firing activity (p < 0.05) was small (9% of DA neurons, 10% of Non-DA neurons). Correlation coefficients between firing activity and Δ[DA] and duration were not significantly different from 0 (sign-rank-test, p >0.05).

## Discussion

An extensive literature documents the relationship between VTA neuron activity and nucleus accumbens dopamine release (Cheer et al., 2007; Owesson-White et al., 2012; Patriarchi et al., 2018; Robinson et al., 2003; Sombers et al., 2009). Despite strong evidence that both dopaminergic and non-dopaminergic neurons coordinate their activities to modulate dopamine release down-stream, few studies have observed these phenomena simultaneously. Thus, the precise relationship between VTA neuron activity and downstream release has not been fully characterized. To investigate how VTA neuronal ensembles influence dopamine release, we simultaneously recorded neural activity in the VTA and pharmacologically induced dopamine release in the NAc of anesthetized rats. We found that the activities of both dopaminergic and non-dopaminergic neurons were, on average, concomitant with the onset of NAc dopamine release. Even so, the precise timing of each neuron’s response (*e*.*g*., time of peak firing) was notably heterogenous, with subsets of dopaminergic and non-dopaminergic neurons responding prior to release, at release onset, or during release. Finally, we observed that transient-to-transient variation in the firing activity of dopaminergic, but not non-dopaminergic neurons, was positively correlated with the magnitude of each dopamine transient. Taken together, these data suggest that dopaminergic and non-dopaminergic neurons in the VTA can work cooperatively to sculpt the timing and magnitude of release.

### Validation of VTA-evoked dopamine release events

Phasic dopamine release events were consistently observed 5 - 10 minutes following eticlopride injection, and these events were eliminated after baclofen injection into the VTA (**Figure 1E-G**). Though the induced dopamine release events are dissimilar in frequency from those that occur in behaving animals, they are comparable in amplitude (see Hamid et al., 2015 for examples of dopamine release in behaving animals). Patterns of VTA activity that evoke dopamine release with this drug cocktail should therefore be similar to neural activity observed in behaving animals.

The elimination of NAc dopamine transients following intra-VTA baclofen injection shows that VTA neuron activity is required for NAc dopamine release in this paradigm. However, this does not imply that dopamine release cannot occur independently of VTA neuron firing, especially considering recent evidence for intra-NAc modulation of release (Chuhma et al., 2014; Threlfell et al., 2012; Zhang et al., 2018). Rather, it shows that D2 receptor antagonism mediates dopamine release at the level of the cell body, and blocking somatodendritic inhibition increases dopamine release in the NAc (Ford, 2014).

### Individual dopaminergic and non-dopaminergic neurons respond at distinct phases of the transient release event, suggesting cooperation in shaping the time course of release

Analysis of the peri-event responses of dopaminergic and non-dopaminergic neurons indicated that maximal responses were, on average, centered around the time of NAc dopamine transient onset and that firing activity was higher surrounding transient onset relative to transient peak (**Figure 3F**). Even so, the precise timing of each individual neuron’s response (*e*.*g*., time of maximal firing) was notably heterogenous, with subsets of neurons responding prior to release onset, at release onset, or during release (**Figure 3A-D**), and dopaminergic and non-dopaminergic neurons responded with both decreases and increases in activity at the time of release onset (**Figure 3H**). Furthermore, there was no difference between dopaminergic and non-dopaminergic neurons with respect to timing relative to the onset of the release event (**Figure 3I**). Taken together, these observations run counter to our original hypothesis that non-dopaminergic inhibitory neurons would reduce activity near the time of release onset to disinhibit dopaminergic neurons. Instead, we found no difference in the number of neurons with increasing or decreasing patterns of activity at the time of release onset (**Figure 3H**). It is worth noting that we were only able to simultaneously record from small subsets of VTA neurons. Consequently, we did not have the statistical power to determine if short-latency interactions between dopaminergic and non-dopaminergic neurons (*e*.*g*., via cross-correlation) suggested disinhibition. Future studies using high-density recording methods could address this.

Our data suggests that variation in the tuning and timing of VTA neuron activity to release may serve to shape the time-course of release. This is further supported by the analysis of spike-triggered responses to the rate of change (estimated first derivative) of dopamine concentration (**Figure 4**). Analysis of these data again indicated that dopamine levels increased, on average, after dopamine and non-dopamine neuron action potentials (**Figure 4E**); however, inspection of the individual STA responses indicated that subsets of VTA neurons responded to rising and falling levels of dopamine release (**Figure 4C, D**).

Finally, while dopaminergic and non-dopaminergic neurons responded heterogeneously to the phase of release, the magnitude of downstream release was positively correlated with transient-to-transient variations in dopaminergic neuron but not non-dopaminergic neuron firing activity (**Figure 5**). Thus, while VTA non-dopaminergic neurons may work with dopaminergic neurons to sculpt the time course of release, the magnitude of release is, perhaps as expected, best correlated with dopamine neuron firing rates.

### Dopaminergic and non-dopaminergic neurons were tonically active

Dopaminergic neurons have tonic and phasic modes of activity (Grace, 1991; Paladini and Roeper, 2014), and these modes likely contribute to distinct functions. For example, fast, phasic release is associated with expectation-driven learning and attention (Badrinarayan et al., 2012; Brischoux et al., 2009; de Jong et al., 2019; Fiorillo et al., 2013; Kobayashi and Schultz, 2014; Matsumoto and Hikosaka, 2009; Mirenowicz and Schultz, 1996; Schultz, 2016c; Schultz et al., 2015, 1993) and working memory (Matsumoto and Takada, 2013). In contrast, tonic activity has been associated with slow-changes in ‘baseline’ dopamine concentration, locomotor activity (Freed and Yamamoto, 1985), motivation (Salamone and Correa, 2012), and response vigor (Niv et al., 2007).

The present experiment investigated fast ‘phasic’ dopamine transients in the NAc. Consequently, we anticipated that dopaminergic neurons would exhibit more burst firing than non-dopaminergic neurons at the onset of each dopamine release event. To address this, a firing-rate independent measure of burst and tonic activity was used to analyze dopaminergic and non-dopaminergic neurons (Shinomoto et al., 2009, 2005). This was important given the considerable variability in firing rates observed across the population of neurons. Indeed, classic mechanisms of burst detection (*e*.*g*., 80/160 ms ISI (Grace and Grace, 1984)) are likely ineffective for dopaminergic neurons under the pharmacological manipulation used here. This analysis indicated that both dopaminergic and non-dopaminergic neurons, on average, expressed tonic activity (local variance < 1; **Figure 3G**), with local variance values being lower for dopaminergic cells. Furthermore, the local variance measure did not change from the time of transient onset to the time of peak release (**Supplemental Figure 4**). Though burst firing is clearly a critical mechanism involved in eliciting phasic dopamine release in behaving animals, these data suggest that increased burst firing may not be essential for phasic dopamine release in the experimental design presented here (see *Limitations* below). For example, eticlopride acts to block activity at D2 receptors, sending dopaminergic neurons into a hyperexcitable state that might limit the ability of cells to burst.

### Limitations

This experiment was designed to investigate VTA firing patterns associated with dopamine release events. Studying such phenomena in awake behaving animals is always preferred, but, in this instance, practical limitations necessitated the use of anesthesia. Carbon-fiber microelectrodes are notoriously fragile and loading them into micro-drives with movable electrode arrays is non-trivial, leading to a low implantation success rate. Behaving experiments would require many more animals, and far more time and materials to acquire similar data.

Although pharmacological manipulation generated frequent, high-magnitude dopamine release events, the cellular and molecular mechanisms that elicit these events were surely altered (Ford, 2014), and comparisons between the data shown here and that of awake, behaving animals should be made with an abundance of caution. D2 antagonists like eticlopride alter normal dopaminergic cell firing by acting on D2 autoreceptors to increase baseline firing (Pucak and Grace, 1994). Additionally, although isoflurane does not appear to alter electrically-stimulated dopamine release (Brodnik and España, 2015), its action on gamma-aminobutyric acid type A (GABA_A_) receptors, glycine receptors, and glutamate receptors likely alter dopaminergic cell firing directly (*e*.*g*., glutamate receptors) or indirectly via non-dopaminergic neurons (*e*.*g*., GABA_A_) (Diao et al., 2014). Consequently, caution is warranted when comparing these results to unanesthetized preparations.

Finally, the method of measuring dopamine release used here might contribute to the temporal heterogeneity observed. Although FSCV is relatively fast, the 5 Hz sampling rate needed to optimize the time between scans for electrophysiological recording decreased the fidelity of the association between the two signals. However, dopamine release events are much longer under the influence of GBR-12909 due to blocked reuptake, and faster events are less prevalent in these data. Future studies using cell-type specific recording methods and tools capable of measuring dopamine release with faster sampling rates (*e*.*g*., dLight, GRAB-DA, (Patriarchi et al., 2018; Sun et al., 2018)) will be needed to examine the role that these heterogeneous signals play in the generation of dopamine release.

### Conclusions

Here, we recorded the firing activities of VTA neurons while simultaneously monitoring pharmacologically induced phasic dopamine release in the ventral striatum (nucleus accumbens core) of anesthetized rats. Although, on average, VTA neuron activity increased at the time of release onset and decreased at the time of peak release, the tuning of individual VTA neurons, regardless of type, was notably heterogenous. Indeed, subsets of neurons responded prior to release, at release onset, and during release. Other neurons were notably silent during release but active otherwise. These data suggest that populations of VTA neurons become active at distinct phases of a dopamine release event and to differing rates of release. Furthermore, the firing activities of dopaminergic but not non-dopaminergic neurons were positively correlated with release magnitude, suggesting that while both cell types may be involved in shaping the time course of release, dopaminergic neurons directly regulate release magnitude. Taken together, the temporal and directional heterogeneity of responses suggests that dopaminergic and non-dopaminergic neurons are not just firing in a ‘population-burst’ fashion, but rather are coordinating their activity to modulate dopamine release events as they are happening, allowing for flexible tuning of dopamine release on a moment-to-moment basis.

## Materials and Methods

### Surgical procedure and implantation

Male Sprague-Dawley rats (n = 22; 350 - 500 g) were anesthetized using 1 - 3% isoflurane in oxygen with a constant 1.5 L/min flow rate. Isoflurane was lowered after induction and implantation until the animal’s breathing rate stabilized at ∼60 breaths per minute (1% - 1.5 % isoflurane mixed with oxygen). Isoflurane doses were kept constant to minimize anesthesia-associated changes in dopamine release. Craniotomies were drilled to target the core of the NAc (AP: 1.4 mm, ML: 1.4 mm from bregma), the VTA (AP: -5.2 to -5.5 mm, ML: 0.4 to 1.0 mm) (Paxinos and Watson, 2009), and the caudal or “tail” component of the VTA (tVTA; AP: -6.5 to -7.5 mm, ML: 0.4 to 1.0 mm). A carbon-fiber microelectrode was then lowered into the NAc core (DV: -6.2 mm to -6.7 mm) and custom-made microelectrode arrays were lowered into the VTA (−7.2 mm to -8.8 mm) and tVTA (DV: -7.2 mm to -8.8 mm). Data acquired from the tVTA were not analyzed in this study. A schematic of the implantation arrangement is in **Figure 1A**. Experiments lasted from 4 – 7 hours from the onset of anesthesia to euthanasia. All procedures were performed in accordance with National Institutes of Health guidelines for laboratory animals under protocols approved by the University of Arizona Institutional Animal Care and Use Committee.

### Pharmacological induction of transient dopamine release and increased dopamine cell firing

Transient dopamine release events were induced pharmacologically using a dopamine transporter (DAT) inhibitor, GBR-12909 dihydrochloride (17 mg/kg at 5 mg/ml; Sigma Aldrich, St. Louis, MO), and a D2 dopamine receptor (D2R) antagonist, S-(−)-Eticlopride hydrochloride (0.75 mg/kg at 5 mg/ml; Sigma Aldrich, St. Louis, MO). GBR-12909 increases extracellular dopamine available for measurement by FSCV by blocking dopamine reuptake, and eticlopride (D2R antagonist) decreases autoreceptor-mediated inhibition of dopamine neurons to increase firing of dopamine neurons (Ford, 2014; Herrik et al., 2010).

A single recording session began with the implantation of the carbon-fiber microelectrode and the microelectrode array followed by *i*.*p*. injection of GBR-12909 (**Figure 1A,B**). Because uptake is slower for GBR-12909 than eticlopride, we waited 20 minutes following injection of GBR-12909 to inject eticlopride (*i*.*p*.). Single-unit and FSCV signals were recorded until transient dopamine release events began to decrease in amplitude. Subsequent ‘booster’ injections of eticlopride were given approximately once every hour until transient dopamine release events were no longer detectable, likely indicating that GBR-12909 was no longer available at effective concentrations.

### Baclofen microinjection

To verify the role of VTA cell firing in driving dopamine release, a GABA_B_ receptor agonist (baclofen) was used to inactivate dopamine cell firing (**Figure 1E-G**). Dopamine release was induced pharmacologically as described in the previous section. Once spontaneous dopamine release events were observed (> 55 nM), a 10-minute baseline was recorded and an intra-VTA injection of baclofen (0.2 mg in 0.5 µL saline) or saline (0.5 µL) was administered at a rate of 0.5 µL/min using a Hamilton 2.5 µL syringe (model 62 RN, part no. 7632-01) and small hub needle (part no. 7803-07, Hamilton Company, Reno, NV). Data were then recorded for ∼ 20 minutes post-injection.

### Electrophysiology

Microelectrode arrays were manufactured in-house using Frederick Haer sharp-electrodes (UE(KG4), Bowdoin, ME) or stereotrodes (McNaughton et al., 1983). Robust firing activity in the red nucleus was used as a landmark when lowering electrodes to improve targeting of the VTA as this nucleus contains high-amplitude fast-spiking neurons and lies 1 mm above the VTA. After identifying cells of the red nucleus, the array was lowered until single-unit spikes with characteristic features of dopamine neurons were detected (Ungless and Grace, 2012). Data were collected on a Windows PC using an Intan RHD2000 USB interface board, an RHD2132 head stage, and RHD2000 interface software version 1.5.2 (Intan Technologies, Los Angeles, CA). Data were sampled at 30 kHz and digitized with 16-bit resolution. Single-unit activity (spikes) from individual neurons were identified by spike-sorting data from the continuous local-field signal (Butterworth band-pass filter 300 - 6000 Hz) using Spike2 software (version 8.12; Cambridge Electronic Design, Milton, Cambridge, England).

### Fast-scan cyclic voltammetry

Carbon-fiber microelectrodes were manufactured as previously described (Hill et al., 2017). Briefly, AS4 carbon fibers (Hexel, Stamford, CT) were aspirated into glass capillaries (0.68 mm I.D.; A-M Systems, Inc., Sequim, WA) and a DKI - 700C Vertical Pipette Puller (David Kopf Instruments, Tujunga, CA) was used to pull electrodes to a point and seal the glass around the carbon fiber. Fibers were cut 70 - 80 mm in length from the glass seal. PEDOT: Nafion was electrodeposited onto the fiber surface following procedures described by Vreeland *et al*. (2014). Ag/AgCl reference electrodes were made by soaking silver wire (0.5 mm diameter, Sigma-Aldrich, St. Louis, MO) in chlorine bleach for 24 h. FSCV measurements were made using custom hardware and software developed by the Heien laboratory and designed to interface with electrophysiological recordings. After implanting a carbon fiber into the NAc, a triangle waveform (scan) was applied at 5 Hz (waveform duration: 8.5 ms at 400 V/s).

### Identification of putative dopaminergic neurons

Neurons were classified as putative dopaminergic or non-dopaminergic neurons using methods similar to Roesch *et al*. (2007). Hierarchical agglomerative clustering based on the waveform amplitude ratio and spike half-width (n clusters = 2, clusterdata function in Matlab) was used to identify an initial segmentation of dopamine and non-dopamine clusters (**Figure 1C**). Outliers were then determined as in Roesch et al. (2007) as cells with waveform parameters that were > 3 standard deviations from each cluster mean. ‘Borderline’ neurons that approached the linear discriminant line separating the classes were eliminated to reduce the misclassification rate (‘+’ in **Figure 1C**). These cells were determined as cells with Fisher’s linear discriminant values that exceeded the 95^th^ percentile or fell below the 5^th^ percentile of each cluster. Neurons were also excluded from analysis if there was clear drift in waveform amplitude through the duration of the recording session. Sessions were excluded from analysis if < 12 transients occurred.

### Histology

Upon the termination of the recording, electrolytic lesions were made at each recording site and animals were sacrificed using 0.35 ml euthanasia solution (390 mg/ml pentobarbital sodium and 50 mg/ml phenytoin sodium; Vetone, Boise, ID) and perfused with 4% paraformaldehyde in phosphate-buffered saline. Brains were extracted and sectioned, and tissue was stained using cresyl violet. Coordinates of electrodes were verified by visual inspection of sectioned tissue. Electrode placements in the VTA was verified histologically, with the most rostral and most caudal electrode lesions sites shown in **Supplemental Figure 1**.

### Data analysis

Data were analyzed using Matlab (Mathworks, Natick, MA). The onsets, terminations, and peaks of dopamine release events were manually scored from the FSCV signal. Change in dopamine release was categorized as a transient dopamine release event if an increase > 1 nA was observed (∼ 55 nM). Recording sessions with < 12 transients were not analyzed. **Burst detection:** Firing rates of putative dopamine neurons were higher than typically reported which was likely due to the application of the D2R antagonist (Herrik et al., 2010) as blocking D2 receptors can increase neuronal excitability (Ohara et al., 1988). As a result, methods of burst detection that rely on fixed inter-spike interval thresholds (*e*.*g*., 80 ms as in (Grace and Bunney, 1984)) will result in false positives. To address this, we used the local variance (LV) measure of bursting (Shinomoto et al., 2009, 2005). This method quantifies burst-like firing activity as the degree to which local changes in inter-spike interval sequences fall below or rise above activity predicted by a Poisson inter-spike interval distribution. LV values < 1 suggest tonic firing, values near 1 suggest Poisson firing, and values > 1 suggest burst-like firing. Uniquely, the local variance measure is robust to non-stationary changes in firing rate and can detect burst-like firing in neurons with high mean firing rates. **Spike Triggered Averages:** Spike-triggered averages (STAs) (Dayan and Abbott, 2001; de Boer and Kuyper, 1968; Schwartz et al., 2006) were used to analyze the time course of dopamine release surrounding individual action potentials. STAs were created by first finding the peri-event normalized dopamine concentrations in a ± 1.5 sec window surrounding each action potential, and then averaging these peri-event responses to determine the mean concentration preceding and following each action potential. This approach can be biased by non-stationarities in the statistics of the action potentials or dopamine release. As a result, a ‘jitter’-averaged STA was generated by randomly jittering the inter-spike intervals of each neuron (drawn from a uniform distribution) in the ± 3 second window surrounding each release event. The jitter-averaged STA was then subtracted from the original STA to reduce the influence of slow changes in rate or release. STAs of Δ[DA] (change in [DA]) were calculated as described above but using the estimated first derivative of [DA] (Matlab: diff function applied to the [DA] signal). Thus, STAs of Δ[DA] represent the direction of dopamine concentration change (*e*.*g*., increasing or decreasing) surrounding each action potential. We also investigated the acceleration of the dopamine signal (if the rate of change of the signal was increasing or decreasing) which was computed as the estimated first derivative of Δ[DA].

## Acknowledgments

We thank Dr. Carol Barnes for the generous donation of electrodes.

## Author contributions

Dan Hill: data collection, data analysis, wrote manuscript

Zachary G. Olson: data collection

Mitchell J. Bartlett: data collection (histology)

Torsten Falk: wrote manuscript

Michael L. Heien: experiment design, data analysis, wrote manuscript

Stephen L. Cowen: experiment design, data analysis, wrote manuscript

## References

Badrinarayan A, Wescott SA, Vander Weele CM, Saunders BT, Couturier BE, Maren S, Aragona BJ. 2012. Aversive Stimuli Differentially Modulate Real-Time Dopamine Transmission Dynamics within the Nucleus Accumbens Core and Shell. Journal of Neuroscience 32:15779–15790. doi:10.1523/JNEUROSCI.3557-12.2012

Bass CE, Grinevich VP, Vance ZB, Sullivan RP, Bonin KD, Budygin EA. 2010. Optogenetic control of striatal dopamine release in rats. Journal of Neurochemistry 114:1344–1352. doi:10.1111/j.1471-4159.2010.06850.x

Bourdy R, Barrot M. 2012. A new control center for dopaminergic systems: Pulling the VTA by the tail. Trends in Neurosciences 35:681–690. doi:10.1016/j.tins.2012.06.007

Brischoux F, Chakraborty S, Brierley DI, Ungless MA. 2009. Phasic excitation of dopamine neurons in ventral VTA by noxious stimuli. Proceedings of the National Academy of Sciences 106:4894–4899. doi:10.1073/pnas.0811507106

Brodnik ZD, España RA. 2015. Dopamine uptake dynamics are preserved under isoflurane anesthesia. Neuroscience Letters 606:129–134. doi:10.1016/j.neulet.2015.08.046

Chuhma N, Mingote S, Moore H, Rayport S. 2014. Dopamine neurons control striatal cholinergic neurons via regionally heterogeneous dopamine and glutamate signaling. Neuron 81:901–912. doi:10.1016/j.neuron.2013.12.027

Da Silva JA, Tecuapetla F, Paixão V, Costa RM. 2018. Dopamine neuron activity before action initiation gates and invigorates future movements. Nature 554:244–248. doi:10.1038/nature25457

Dayan P, Abbott LF. 2001. Theoretical neuroscience : computational and mathematical modeling of neural systems. Cambridge, Mass: Massachusetts Institute of Technology Press.

de Boer E, Kuyper P. 1968. Triggered Correlation. IEEE Transactions on Biomedical Engineering BME-15:169–179. doi:10.1109/TBME.1968.4502561

de Jong JW, Afjei SA, Pollak Dorocic I, Peck JR, Liu C, Kim CK, Tian L, Deisseroth K, Lammel S. 2019. A Neural Circuit Mechanism for Encoding Aversive Stimuli in the Mesolimbic Dopamine System. Neuron 101:133-151.e7. doi:10.1016/j.neuron.2018.11.005

Diao S, Ni J, Shi X, Liu P, Xia W. 2014. Mechanisms of action of general anesthetics. Frontiers in Bioscience-Landmark 19:747–757. doi:10.2741/4241

Engelhard B, Finkelstein J, Cox J, Fleming W, Jang HJ, Ornelas S, Koay SA, Thiberge SY, Daw ND, Tank DW, Witten IB. 2019. Specialized coding of sensory, motor and cognitive variables in VTA dopamine neurons. Nature 570:509–513. doi:10.1038/s41586-019-1261-9

Eshel N, Tian J, Bukwich M, Uchida N. 2016. Dopamine neurons share common response function for reward prediction error. Nature Neuroscience 19:479–486. doi:10.1038/nn.4239

Fiorillo CD, Song MR, Yun SR. 2013. Multiphasic temporal dynamics in responses of midbrain dopamine neurons to appetitive and aversive stimuli. Journal of Neuroscience 33:4710–4725. doi:10.1523/JNEUROSCI.3883-12.2013

Ford CP. 2014. The role of D2-autoreceptors in regulating dopamine neuron activity and transmission. Neuroscience. doi:10.1016/j.neuroscience.2014.01.025

Freed CR, Yamamoto BK. 1985. Regional brain dopamine metabolism: a marker for the speed, direction, and posture of moving animals. Science (New York, NY) 229:62–5.

Grace A, Grace A. 1984. Grace AA, Bunney BS. The control of firing pattern in nigral dopamine nurons : burst firing. J Neurosci 4 : 2877-2890 2877–2890.

Grace AA. 1991. Phasic versus tonic dopamine release and the modulation of dopamine system responsivity: a hypothesis for the etiology of schizophrenia. Neuroscience 41:1–24. doi:10.1016/0306-4522(91)90196-u

Grace AA, Bunney BS. 1984. The control of firing pattern in nigral dopamine neurons: burst firing. The Journal of neuroscience : the official journal of the Society for Neuroscience 4:2877–90.

Hamid AA, Pettibone JR, Mabrouk OS, Hetrick VL, Schmidt R, Vander Weele CM, Kennedy RT, Aragona BJ, Berke JD. 2015. Mesolimbic dopamine signals the value of work. Nature Neuroscience 19:117–126. doi:10.1038/nn.4173

Hart AS, Rutledge RB, Glimcher PW, Phillips PEM. 2014. Phasic dopamine release in the rat nucleus accumbens symmetrically encodes a reward prediction error term. The Journal of neuroscience : the official journal of the Society for Neuroscience 34:698–704. doi:10.1523/JNEUROSCI.2489-13.2014

Herrik KF, Christophersen P, Shepard PD. 2010. Pharmacological Modulation of the Gating Properties of Small Conductance Ca2+ Activated K+ Channels Alters the Firing Pattern of Dopamine Neurons In Vivo. Journal of Neurophysiology 104:1726–1735. doi:10.1152/jn.01126.2009.

Hill DF, Parent KL, Atcherley CW, Cowen SL, Heien ML. 2017. Differential release of dopamine in the nucleus accumbens evoked by low-versus high-frequency medial prefrontal cortex stimulation. Brain stimulation 11:426–434. doi:10.1016/j.brs.2017.11.010

Howe MW, Dombeck DA. 2016. Rapid signalling in distinct dopaminergic axons during locomotion and reward. Nature 535:505–510. doi:10.1038/nature18942

Jackson ME, Frost AS, Moghaddam B. 2001. Stimulation of prefrontal cortex at physiologically relevant frequencies inhibits dopamine release in the nucleus accumbens. Journal of neurochemistry 78:920–3.

Jalabert M, Bourdy R, Courtin J, Veinante P, Manzoni OJ, Barrot M, Georges F. 2011. Neuronal circuits underlying acute morphine action on dopamine neurons. Proceedings of the National Academy of Sciences 108:16446–16450. doi:10.1073/pnas.1105418108

Jhou TC, Fields HL, Baxter MG, Saper CB, Holland PC. 2009. The Rostromedial Tegmental Nucleus (RMTg), a GABAergic Afferent to Midbrain Dopamine Neurons, Encodes Aversive Stimuli and Inhibits Motor Responses. Neuron 61:786–800. doi:10.1016/j.neuron.2009.02.001

Kobayashi S, Schultz W. 2014. Reward contexts extend dopamine signals to unrewarded stimuli. Current Biology 24:56–62. doi:10.1016/j.cub.2013.10.061

Kuhr WG, Wightman RM, Rebec G V. 1987. Dopaminergic neurons: simultaneous measurements of dopamine release and single-unit activity during stimulation of the medial forebrain bundle. Brain Research 418:122–128. doi:10.1016/0006-8993(87)90968-1

Lisman JE, Grace AA. 2005. The hippocampal-VTA loop: Controlling the entry of information into long-term memory. Neuron 46:703–713. doi:10.1016/j.neuron.2005.05.002

Lohani S, Martig AK, Underhill SM, DeFrancesco A, Roberts MJ, Rinaman L, Amara S, Moghaddam B. 2018. Burst activation of dopamine neurons produces prolonged post-burst availability of actively released dopamine. Neuropsychopharmacology : official publication of the American College of Neuropsychopharmacology 43:2083–2092. doi:10.1038/s41386-018-0088-7

Lüscher C. 2016. The Emergence of a Circuit Model for Addiction. Annual Review of Neuroscience 39:257–276. doi:10.1146/annurev-neuro-070815-013920

Margolis EB, Lock H, Hjelmstad GO, Fields HL. 2006. The ventral tegmental area revisited: is there an electrophysiological marker for dopaminergic neurons? The Journal of physiology 577:907–924. doi:10.1113/jphysiol.2006.117069

Margolis EB, Toy B, Himmels P, Morales M, Fields HL. 2012. Identification of rat ventral tegmental area GABAergic neurons. PLoS ONE 7. doi:10.1371/journal.pone.0042365

Matsui A, Williams JT. 2011. Opioid-Sensitive GABA Inputs from Rostromedial Tegmental Nucleus Synapse onto Midbrain Dopamine Neurons. J Neurosci 31:17729–17735. doi:10.1523/JNEUROSCI.4570-11.2011

Matsumoto M, Hikosaka O. 2009. Two types of dopamine neuron distinctly convey positive and negative motivational signals. Nature 459:837–841. doi:10.1038/nature08028

Matsumoto M, Takada M. 2013. Distinct Representations of Cognitive and Motivational Signals in Midbrain Dopamine Neurons. Neuron 79:1011–1024. doi:10.1016/j.neuron.2013.07.002

McNaughton BL, O’Keefe J, Barnes CA. 1983. The stereotrode: a new technique for simultaneous isolation of several single units in the central nervous system from multiple unit records. J Neurosci Methods 8:391–7.

Melchior JR, Ferris MJ, Stuber GD, Riddle DR, Jones SR. 2015. Optogenetic versus electrical stimulation of dopamine terminals in the nucleus accumbens reveals local modulation of presynaptic release. Journal of Neurochemistry 134:833–844. doi:10.1111/jnc.13177

Menegas W, Akiti K, Amo R, Uchida N, Watabe-Uchida M. 2018. Dopamine neurons projecting to the posterior striatum reinforce avoidance of threatening stimuli. Nature Neuroscience 21:1421–1430. doi:10.1038/s41593-018-0222-1

Mirenowicz J, Schultz W. 1996. Preferential activation of midbrain dopamine neurons by appetitive rather than aversive stimuli. Nature 379:449–451. doi:10.1038/379449a0

Morales M, Margolis EB. 2017. Ventral tegmental area: cellular heterogeneity, connectivity and behaviour. Nature Reviews Neuroscience 18:73–85. doi:10.1038/nrn.2016.165

Niv Y, Daw ND, Joel D, Dayan P. 2007. Tonic dopamine: opportunity costs and the control of response vigor. Psychopharmacology (Berl) 191:507–520. doi:10.1007/s00213-006-0502-4

Ogasawara T, Nejime M, Takada M, Matsumoto M. 2018. Primate Nigrostriatal Dopamine System Regulates Saccadic Response Inhibition. Neuron 100:1513-1526.e4. doi:10.1016/j.neuron.2018.10.025

Ohara K, Haga K, Berstein G, Haga T, Ichiyama A, Ohara K. 1988. The interaction between D-2 dopamine receptors and GTP-binding proteins. Molecular Pharmacology 33:290–296.

Omelchenko N, Sesack SR. 2009. Ultrastructural analysis of local collaterals of rat ventral tegmental area neurons: GABA phenotype and synapses onto dopamine and GABA cells. Synapse 63:895–906. doi:10.1002/syn.20668

Paladini CA, Roeper J. 2014. Generating bursts (and pauses) in the dopamine midbrain neurons. Neuroscience 282:109–121. doi:10.1016/j.neuroscience.2014.07.032

Parent KL, Hill DF, Crown LM, Wiegand J-P, Gies KF, Miller MA, Atcherley CW, Heien ML, Cowen SL. 2017. Platform to Enable Combined Measurement of Dopamine and Neural Activity. Analytical chemistry 89:2790–2799. doi:10.1021/acs.analchem.6b03642

Patriarchi T, Cho JR, Merten K, Howe MW, Marley A, Xiong WH, Folk RW, Broussard GJ, Liang R, Jang MJ, Zhong H, Dombeck D, von Zastrow M, Nimmerjahn A, Gradinaru V, Williams JT, Tian L. 2018. Ultrafast neuronal imaging of dopamine dynamics with designed genetically encoded sensors. Science 360. doi:10.1126/science.aat4422

Paxinos G, Watson C. 2009. The Rat Brain in Stereotaxic Coordinates; Compact, Sixth Edition. Elsevier.

Polter AM, Barcomb K, Tsuda AC, Kauer JA. 2018. Synaptic function and plasticity in identified inhibitory inputs onto VTA dopamine neurons. European Journal of Neuroscience 47:1208–1218. doi:10.1111/ejn.13879

Pucak ML, Grace AA. 1994. Evidence that systemically administered dopamine antagonists activate dopamine neuron firing primarily by blockade of somatodendritic autoreceptors., Journal of Pharmacology and Experimental Therapeutics.

Redgrave P, Gurney K. 2006. The short-latency dopamine signal: a role in discovering novel actions? Nature reviews Neuroscience 7:967–975. doi:10.1038/nrn2022

Roesch MR, Calu DJ, Schoenbaum G. 2007. Dopamine neurons encode the better option in rats deciding between differently delayed or sized rewards. Nature Neuroscience 10:1615–1624. doi:10.1038/nn2013

Root DH, Estrin DJ, Morales M. 2018. Aversion or Salience Signaling by Ventral Tegmental Area Glutamate Neurons. iScience 2:51–62. doi:10.1016/j.isci.2018.03.008

Salamone JD, Correa M. 2012. The Mysterious Motivational Functions of Mesolimbic Dopamine. Neuron 76:470–485. doi:10.1016/j.neuron.2012.10.021

Schultz W. 2016a. Dopamine reward prediction-error signalling: a two-component response. Nature Reviews 17.

Schultz W. 2016b. Dopamine reward prediction-error signalling: a two-component response. Nature Reviews 17.

Schultz W. 2016c. Dopamine reward prediction error coding. Dialogues in Clinical Neuroscience 18:23–32. doi:10.1523/jneurosci.1950-18.2018

Schultz W. 2015. Neuronal Reward and Decision Signals: From Theories to Data. Physiological Reviews 95:853–951. doi:10.1152/physrev.00023.2014

Schultz W, Apicella P, Ljungberg T. 1993. Responses of monkey dopamine neurons to reward and conditioned stimuli during successive steps of learning a delayed response task. Journal of Neuroscience 13:900–913. doi:10.1523/jneurosci.13-03-00900.1993

Schultz W, Carelli RM, Wightman RM. 2015. Phasic dopamine signals: From subjective reward value to formal economic utility. Current Opinion in Behavioral Sciences 5:147–154. doi:10.1016/j.cobeha.2015.09.006

Schultz W, Stauffer WR, Lak A. 2017. The phasic dopamine signal maturing: from reward via behavioural activation to formal economic utility. Current Opinion in Neurobiology, Neurobiology of Learning and Plasticity 43:139–148. doi:10.1016/j.conb.2017.03.013

Schwartz O, Pillow JW, Rust NC, Simoncelli EP. 2006. Spike-triggered neural characterization. Journal of Vision 6:484–507. doi:10.1167/6.4.13

Shinomoto S, Kim H, Shimokawa T, Matsuno N, Funahashi S, Shima K, Fujita I, Tamura H, Doi T, Kawano K, Inaba N, Fukushima K, Kurkin S, Kurata K, Taira M, Tsutsui K-I, Komatsu H, Ogawa T, Koida K, Tanji J, Toyama K. 2009. Relating Neuronal Firing Patterns to Functional Differentiation of Cerebral Cortex. PLoS Computational Biology 5:e1000433. doi:10.1371/journal.pcbi.1000433

Shinomoto S, Miyazaki Y, Tamura H, Fujita I. 2005. Regional and laminar differences in in vivo firing patterns of primate cortical neurons. Journal of Neurophysiology 94:567–575. doi:10.1152/jn.00896.2004

Sombers LA, Beyene M, Carelli RM, Mark Wightman R. 2009. Synaptic Overflow of Dopamine in the Nucleus Accumbens Arises from Neuronal Activity in the Ventral Tegmental Area. Journal of Neuroscience 29:1735–1742. doi:10.1523/JNEUROSCI.5562-08.2009

Stefani MR, Moghaddam B. 2006. Rule learning and reward contingency are associated with dissociable patterns of dopamine activation in the rat prefrontal cortex, nucleus accumbens, and dorsal striatum. Journal of neuroscience 26:8810–8.

Sugam JA, Day JJ, Wightman RM, Carelli RM. 2012. Phasic nucleus accumbens dopamine encodes risk-based decision-making behavior. Biol Psychiat 71:199–205. doi:10.1016/j.biopsych.2011.09.029

Sun F, Zeng J, Jing M, Zhou J, Feng J, Owen SF, Luo Y, Li F, Wang H, Yamaguchi T, Yong Z, Gao Y, Peng W, Wang L, Zhang S, Du J, Lin D, Xu M, Kreitzer AC, Cui G, Li Y. 2018. A Genetically Encoded Fluorescent Sensor Enables Rapid and Specific Detection of Dopamine in Flies, Fish, and Mice. Cell 174:481-496.e19. doi:10.1016/j.cell.2018.06.042

Tan KR, Yvon C, Turiault M, Mirzabekov JJ, Doehner J, Labouèbe G, Deisseroth K, Tye KM, Lüscher C. 2012. GABA Neurons of the VTA Drive Conditioned Place Aversion. Neuron 73:1173–1183. doi:10.1016/j.neuron.2012.02.015

Threlfell S, Lalic T, Platt NJ, Jennings KA, Deisseroth K, Cragg SJ. 2012. Striatal dopamine release is triggered by synchronized activity in cholinergic interneurons. Neuron 75:58–64. doi:10.1016/j.neuron.2012.04.038

Ungless MA, Grace AA. 2012. Are you or aren’t you? Challenges associated with physiologically identifying dopamine neurons. Trends in Neurosciences 35:422–430. doi:10.1016/j.tins.2012.02.003

van Zessen R, Phillips JL, Budygin EA, Stuber GD. 2012. Activation of VTA GABA Neurons Disrupts Reward Consumption. Neuron 73:1184–1194. doi:10.1016/j.neuron.2012.02.016

Vreeland RF, Laude ND, Lambert SM, Heien ML. 2014. Microwave-plasma dry-etch for fabrication of conducting polymer microelectrodes. Analytical chemistry 86:1385–90. doi:10.1021/ac403363a

Xi Z-X, Stein EA. 1998. Nucleus accumbens dopamine release modulation by mesolimbic GABAA receptors—an in vivo electrochemical study. Brain Research 798:156–165. doi:http://dx.doi.org/10.1016/S0006-8993(98)00406-5

Zhang YF, Reynolds JNJ, Cragg SJ. 2018. Pauses in Cholinergic Interneuron Activity Are Driven by Excitatory Input and Delayed Rectification, with Dopamine Modulation. Neuron 98:918-925.e3. doi:10.1016/j.neuron.2018.04.027

