## Supplemental Figure for "Heterogeneous neuronal activity in the ventral tegmental area coordinates dopamine release in the nucleus accumbens"

Supplemental Figure 1

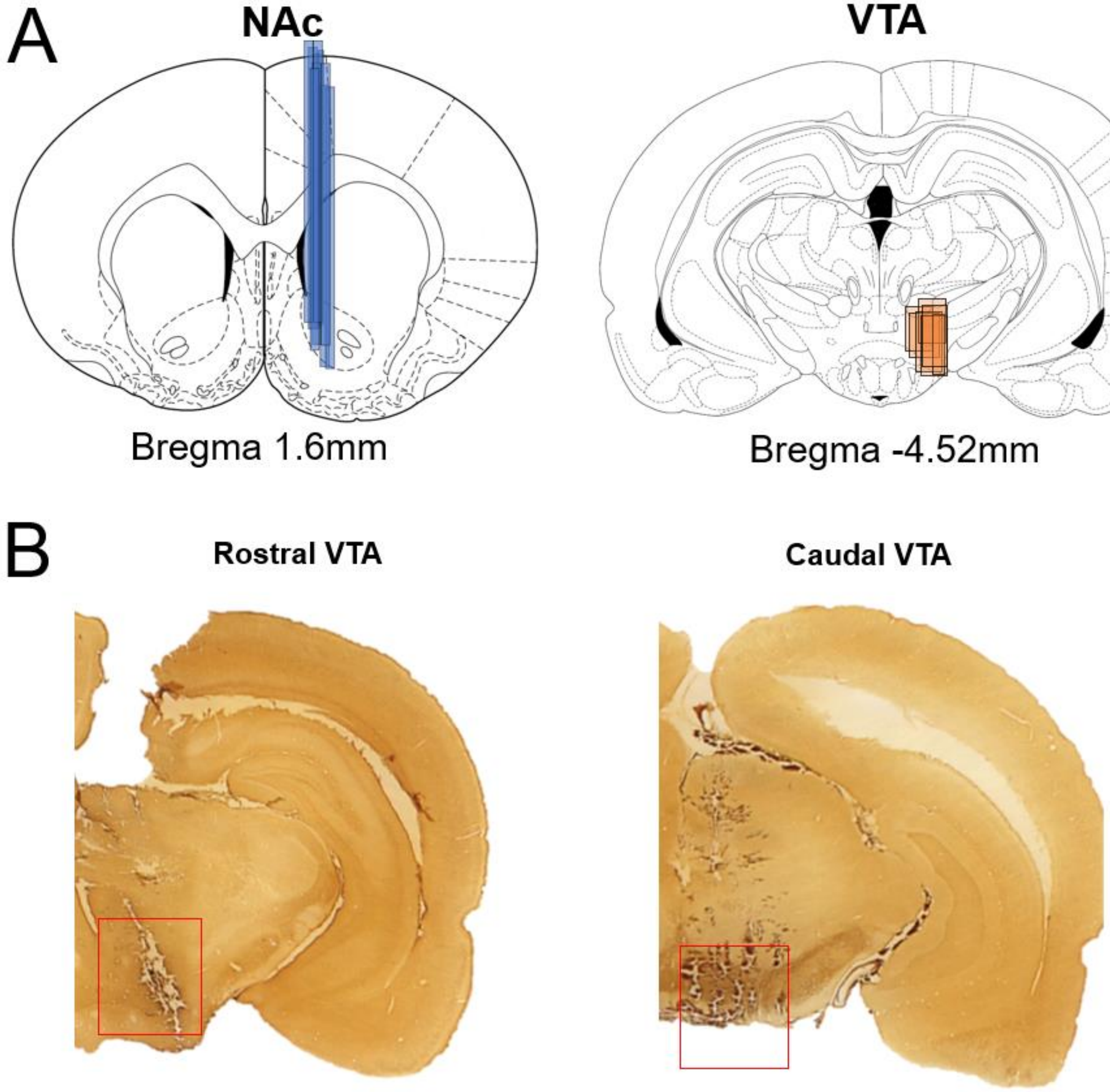


A) Summary of all electrode recording sites in the NAc (FSCV) and VTA (electrophysiology).
B) Histological sections taken from the rostral and caudal regions of the VTA.

Supplemental Figure 2

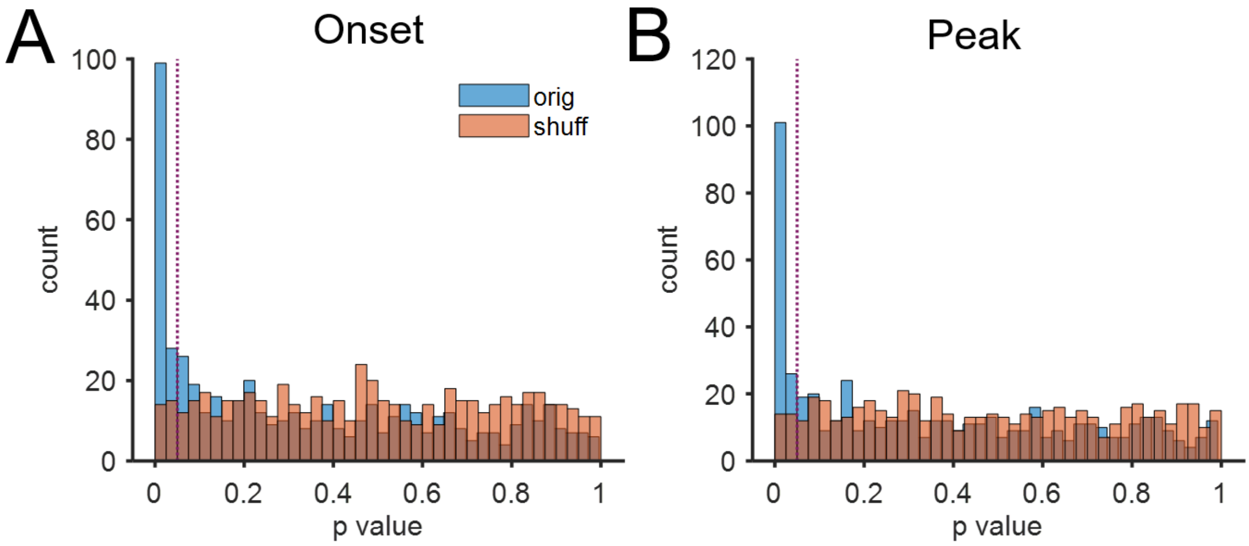


**<u>Assessment of the detection of responsive neurons: Shuffle Comparison</u>**

Our criterion for a selective neurons was $p < 0.05$ for a within-subject Friedman test (non-parametric within-subject test) where the test was applied to a table where rows were the transient ID and columns were the 5 time bins (800 ms bin size) surrounding either release onset or peak release. To validate this criterion, we compared our results with results from applying the same criterion to data produced by shuffling the order of spike counts in each of the five time bins within each trial. The distributions of p values for all neurons surrounding the onset and peak are presented above with blue indicating the p values from the original data and the red indicating p values from the shuffled data. The data indicate that the p-value threshold of 0.05 (vertical line) identified many significant responses above those predicted by chance. We chose the Friedman test to avoid confounds due to assumptions of normality in the data (*e.g.*, due to sparse firing activity and an exponential right tail). In addition, the ratio of true positives to false positives (assuming the shuffle control indicated the false positive detection rate) was 4.4 for the Friedman test but only 3.8 for the within-subject ANOVA for $p < 0.05$.

Supplemental Figure 3

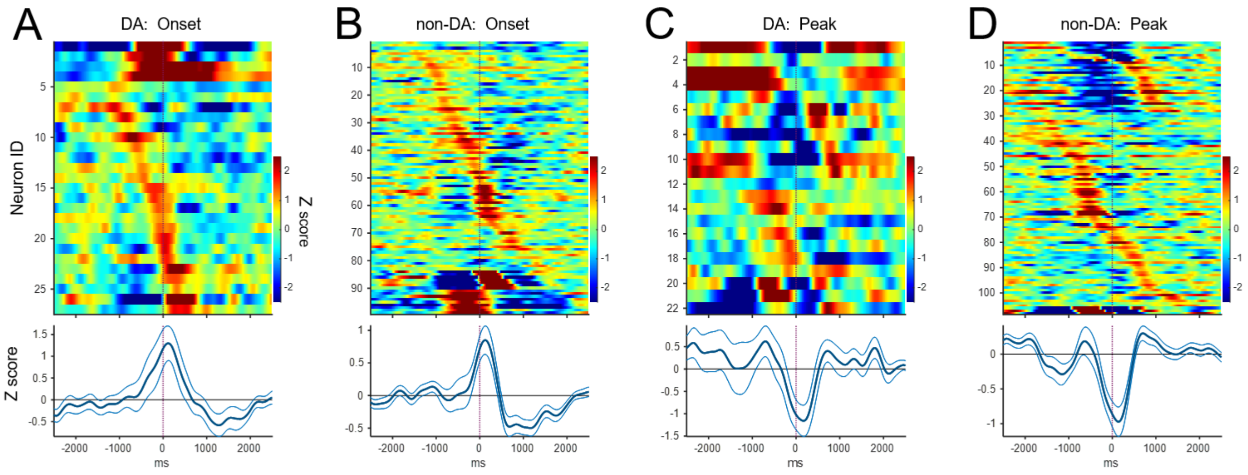


A-D) Average firing rates of all responsive neurons aligned to phasic dopamine release events (data from Figure 3) and sorted by k-means clustering (k = 3). A) Peri-event responses of dopaminergic neurons aligned to the onset of each dopamine transient. Each row indicates the average peri-onset response of a single neuron. Color indicates firing rate in z-scores relative to baseline with baseline defined as activity during the -3 to -4 and +3 to +4 second intervals surrounding the transient onset. Line plot below the color plot indicates the mean response (in z scores, +/- SEM) for all neurons. C & D) As in A & B, but for neurons selective to the time of peak dopamine release. Baseline was defined as -3 to -4 and +3 to +4 second intervals surrounding the transient peak.

Supplemental Figure 4

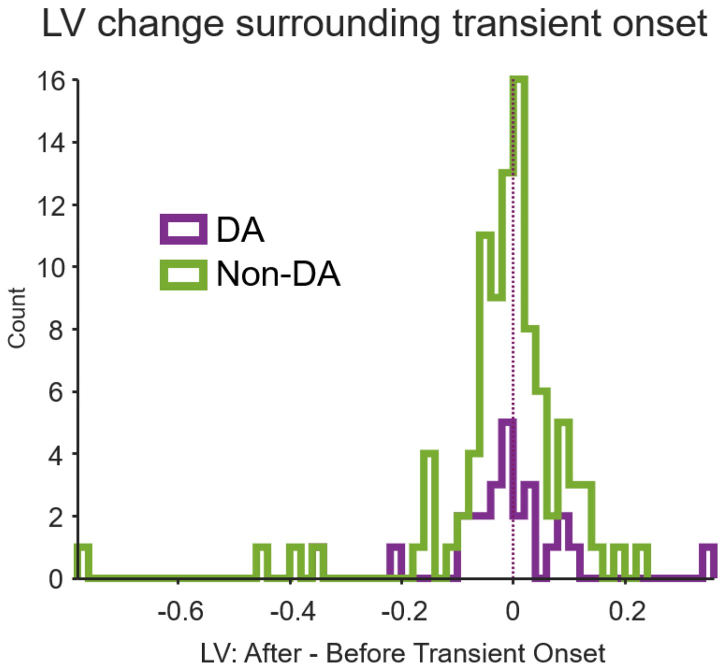


The local variance (LV) measure did not change from the 1 second preceding to the 1 second following transient onset. The x axis indicates the local variance for the 1-second window post transient onset – the 1-second window preceding onset. Positive values indicate an increase in bursting during the transient. The distributions did not differ from zero (pDA = 0.57, pNon-DA = 0.21, paired t-test) for either the DA or non-DA neurons. Only neurons selective to the transient onset were used in this analysis.

Supplemental Figure 5

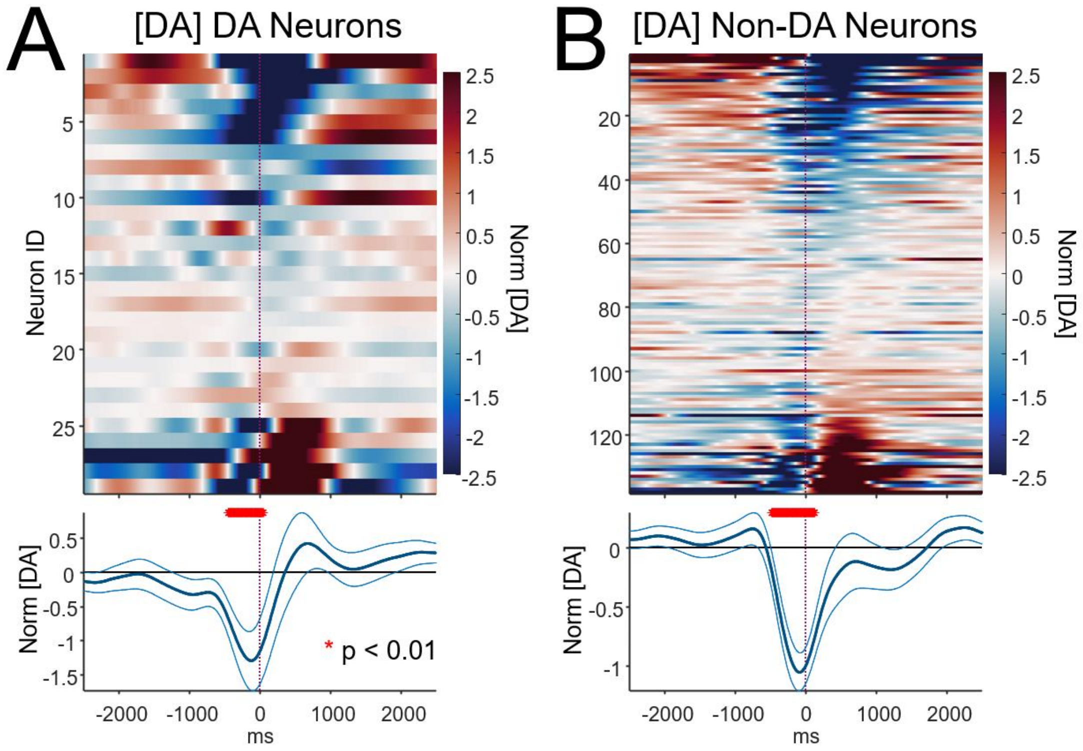


Spike-triggered averages (STAs) of normalized dopamine concentration ([DA]) for dopaminergic and non-dopaminergic neurons. Each row indicates the STA for a single dopaminergic neuron (see Methods for details on STA calculation). Color indicates the z-score of [DA] relative to baseline, with baseline defined as activity during the -3 to -4 and +3 to +4 second intervals surrounding t = 0 (the time of each action potential). For visualization and to highlight the diversity of dopamine dynamics surrounding each action potential, STAs are sorted by k-means cluster groups (k = 3) and, secondarily, by the location of the peak response within each group. The bottom panel shows the mean ±SEM. Red asterisks indicate $p < 0.01$ (Wilcoxon Rank Sum test) relative to zero. The sorted STAs for [DA] for dopaminergic neurons are presented in subplot A and the STAs for non-dopaminergic neurons are presented in B.
